# Deconvolution of HIV-1 Mutational Signatures Reveals Dominant and Donor-Specific APOBEC3-Associated Mutagenesis Across Anatomical Compartments

**DOI:** 10.64898/2026.09.23.753842

**Authors:** Mohadeseh Soleimanpour, Niloofar Haghjoo, Jake D. Lehle, Rachael Springman-Rodriguez, Davey M. Smith, Sara Gianella Weibel, Antoine Chaillon, Diako Ebrahimi

## Abstract

HIV-1 persists across multiple anatomical compartments, yet the mutational processes shaping viral diversity within these reservoirs remain incompletely understood. We investigated HIV-1 mutational signatures across blood, spleen, gut, and brain tissues from 21 people with HIV enrolled in the Last Gift rapid autopsy program. Mutations in single-genome HIV-1 env DNA sequences were quantified by trinucleotide context and deconvolved using non-negative matrix factorization. Two APOBEC3-associated signatures were resolved and together accounted for approximately 50% of all mutations; this contribution remained stable after normalization by the number of potential APOBEC3 target sites. Normalization further revealed a minor signature consistent with methylation-induced cytosine deamination. Mutational patterns were driven primarily by donor identity rather than anatomical compartment. Identical APOBEC3-hypermutated sequences were recovered across brain, gut, and blood, consistent with clonal expansion and trafficking of infected cells. These findings establish APOBEC3 activity as a major contributor to HIV-1 mutagenesis across tissue reservoirs and demonstrate that APOBEC3-associated mutagenesis is strongly donor-specific.

**SIGNIFICANCE:** Despite the ability of antiretroviral therapy to suppress plasma viremia to undetectable levels, HIV persists in diverse anatomical compartments and constitutes the main barrier to viral eradication. This study identifies APOBEC3 enzymes as a dominant source of mutation across anatomical compartments, with a secondary contribution from methylation-associated processes, and shows that their mutational signatures are strongly donor-specific. Establishing APOBEC3 activity as a major, host-driven process clarifies a key source of inter-individual variation in viral evolution, reservoir composition, and responses to therapy. These findings underscore the importance of accounting for donor-level differences in innate antiviral activity when designing strategies to control or eliminate persistent HIV infection.

## INTRODUCTION

In most people with HIV (PWH) HIV-1 replication becomes undetectable in plasma after antiretroviral therapy (ART). However, the virus persists in cellular reservoirs across different compartments including peripheral blood mononuclear cells (PBMCs), gut-associated lymphoid tissue (GALT), the spleen, and the central nervous system (CNS) [1, 2]. These compartments have distinct populations of infected cells, including resting CD4^+^ T cells, tissue-resident macrophages, and microglial cells in the brain [3, 4, 5, 6]. Therefore, viral evolution and selective pressures may differ substantially across anatomical sites, posing a fundamental challenge for HIV control and eradication [7, 8, 9]. Understanding the mutational processes that shape HIV-1 diversity within and between these compartments is critical for informing therapeutic approaches aimed at reservoir elimination.

One of the key mechanisms that directly impacts HIV mutations and genetic diversity, is cytidine deamination by APOBEC3 (Apolipoprotein B mRNA Editing Catalytic Polypeptide-like 3, A3) enzymes [10]. These proteins constitute a major component of the innate immune response against a wide range of viruses including retroviruses such as HIV [11, 12]. The cytoplasmic members of this enzyme family can incorporate into budding virions and, during reverse transcription in newly infected cells, deaminate cytidines to uridines on the minus-strand viral DNA, resulting in characteristic G-to-A hypermutation in the plus-strand coding sequence [13, 14, 15, 16]. Individual A3 family members exhibit distinct sequence context preferences: A3G preferentially introduces GG-to-AG/AA mutations, while other A3 enzymes (e.g. A3D/F/H) preferentially induce GA-to-AA mutations [17, 18, 19, 20]. These changes constitute a substantial fraction of HIV mutations [21, 22].

Importantly, the expression of A3 enzymes is highly cell-type-specific. For instance, T cells and NK cells predominantly express A3G and A3C, and to a lesser extent A3D, A3F, and A3H. By contrast, macrophages and dendritic cells express primarily A3A [23, 24]. Given that different tissues harbor diverse cellular reservoirs for HIV, one would predict that A3-mediated mutational signatures imprinted on the virus should differ across anatomical compartments. To investigate this, we leveraged HIV-1 envelope (*env*) sequences obtained from PWH enrolled in the Last Gift cohort, a unique rapid-autopsy study designed to enable comprehensive multi-compartment analyses of HIV-1 reservoirs [1, 25]. Rapid autopsy procedures enabled extensive tissue collection from each donor, including multiple central nervous system (CNS) regions, gastrointestinal tract sites, lymphoid tissues, peripheral blood, and additional anatomical compartments. This comprehensive tissue sampling provided an unprecedented opportunity to compare viral sequences and their mutational landscapes across anatomical sites within the same person [1, 26].

To accurately identify mutational signatures including those associated with A3 activity, we conducted mutational signature deconvolution using nonnegative matrix factorization (NMF) [26, 27]. This method has been widely used to resolve complex and overlapping mutational signatures in cancer, such as age-associated mutations (SBS1: C-to-T mutations within CG) [28, 29], A3-induced deamination (SBS2/13: C-to-T/G mutations within TCA and TCT) [30, 31], and mutations caused by platinum chemotherapy treatment (SBS31: C-to-T mutations within CCA and CCT) [32, 33]. Applying this approach to HIV is particularly promising because A3-driven mutations are embedded within a broader mutational background generated by other mutational processes, as well as the intrinsic infidelity of reverse transcriptase, which, unlike A3 enzymes, exhibits limited sequence-context dependence [34, 35]. NMF therefore provides a robust framework for disentangling overlapping mutational signatures and quantifying the relative contributions of distinct mutational processes at the level of individual viral sequences.

In this study, we leveraged NMF together with HIV-1 *env* DNA sequences derived from the Last Gift cohort to address three specific questions. First, what are the dominant HIV-1 mutational signatures across anatomical compartments? Here, we applied NMF-based mutational signature deconvolution to identify distinct mutational processes and estimate their relative contributions within individual sequences. Second, what factors shape the distribution of these signatures? Using analysis of variance, we assessed the extent to which mutational signature variations are explained by tissue identity, donor identity, or their interaction, thereby distinguishing local compartments effects from host-specific influences. Third, are viruses carrying similar mutational signatures shared across anatomical compartments? By determining whether similar mutational profiles were detected in multiple tissues, we evaluated evidence for or against the clonal dissemination of infected cells and the connectivity of viral reservoirs across the body.

## RESULTS

### Overview of the HIV-1 Trinucleotide Mutation Dataset

To characterize the mutational signatures observed in HIV-1 *env* DNA, we constructed a trinucleotide mutation matrix from 1,926 HIV-1 *env* single-genome sequences obtained from 21 Last Gift donors (Supplementary Table S1). These sequences were obtained from multiple tissues, including up to 10 CNS (brainstem/basal ganglia [BSG], frontal cortex matter [FCM], hippocampus [HPC], medulla oblongata [MED], occipital cortex [OCC], parietal cortex [PCT], pons [PON], spinal cord cervical [SCC], spinal cord lumbar [SCL], spinal cord thoracic [SCT]) regions, 5 gut (colon/large intestine [CLR], duodenum [DDM], ileum [ILM], jejunum [JJM], rectum [RCT]) regions, spleen, and PBMC per donor. The matrix comprised 1,077 trinucleotide mutation types (columns), with each row representing one of the 1,926 individual sequences (Supplementary Fig S1A). To avoid biases arising from unequal representation of trinucleotide motifs, we normalized mutation counts by the genomic frequency of each trinucleotide motif in each sequence. This ensured unbiased comparisons across sequences, tissues, and donors.

Examination of mutation count distributions across trinucleotides and sequences revealed that a relatively small number of trinucleotide mutation types accounted for most observed mutations, consistent with the action of discrete mutational processes (Supplementary Fig S1B). Sampling depth varied substantially across donors and tissues (Supplementary Fig S2), with particularly limited representation in the spleen, which should be considered when interpreting downstream analyses.

### A3-associated GG-to-AG and GA-to-AA changes are the most frequent mutations

We first quantified the frequency of each mutation type across all donors and anatomical compartments combined. Figure 1A shows the 20 most frequent mutation types based on total mutation counts, with A3-associated G-to-A mutations in GG or GA sequence contexts highlighted in dark color. The five most frequent mutation types were A3-associated, including TGG-to-TAG and AGA-to-AAA, which represent preferred mutation contexts associated with A3G and other A3 enzymes, particularly A3D/F/H, respectively.

**Figure 1.**
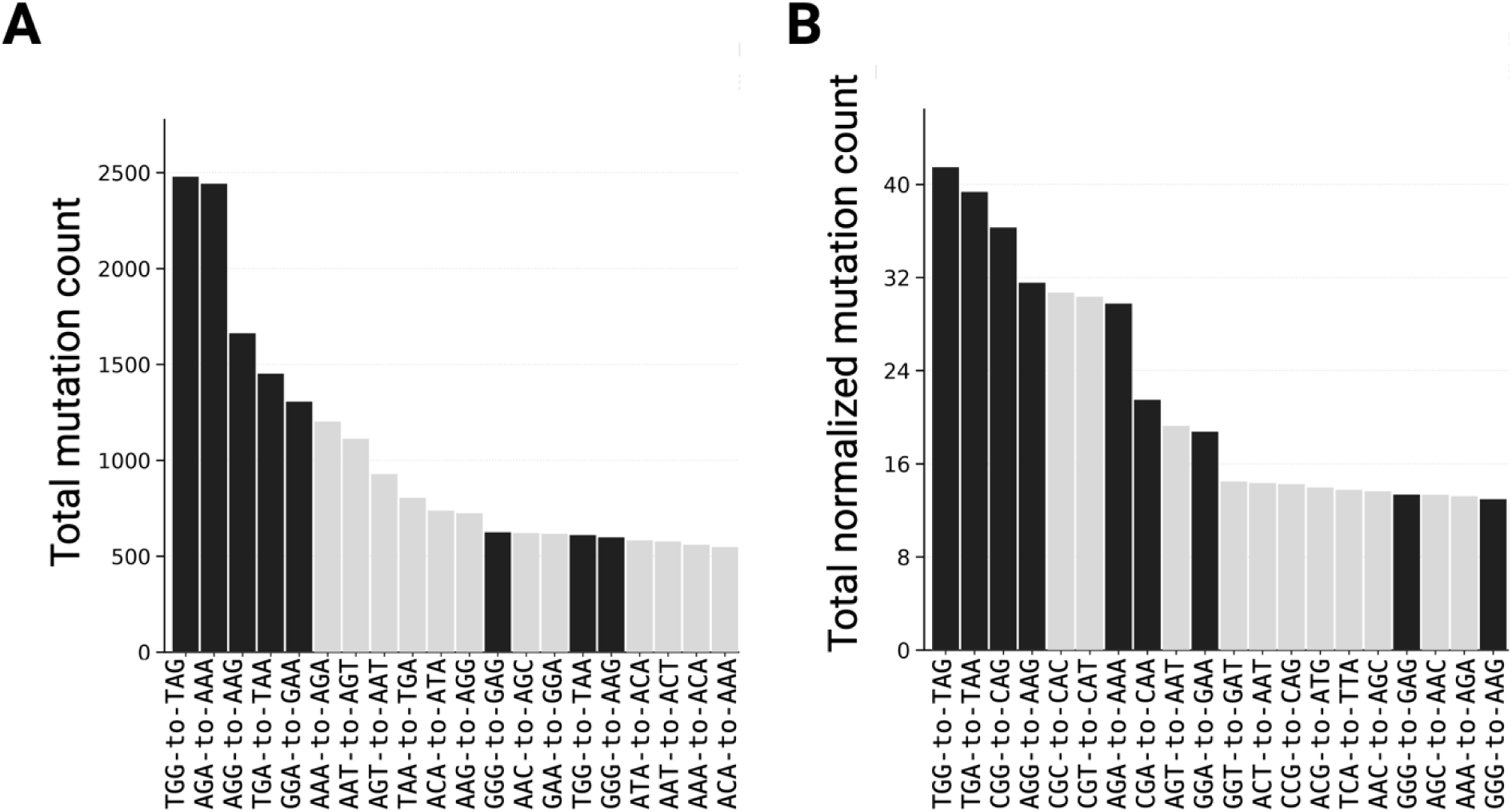
Most frequent HIV-1 mutation types across donors and anatomicalcompartments. (A) Total mutation counts for the 20 most frequent mutation types across all 1,926 sequences. (B) Mutation counts normalized by the total number of corresponding trinucleotide motifs across all sequences. A3-associated G-to-A mutations occurring in GG or GA contexts are shown in black and comprise the most frequent mutation types.

We then normalized the mutation counts for each mutation type by the frequency of its corresponding trinucleotide context across all sequences (Fig. 1B). Importantly, A3-associated G-to-A mutations remained the dominant mutation types after normalization, with four of the five most frequent normalized mutation types corresponding to A3-preferred GG or GA contexts. This persistence indicates that the strong contribution of A3-associated mutagenesis is not simply attributable to differences in the abundance of its target sequences. Normalization also increased the relative frequency of CpG-containing mutations (e.g. CGC-to-CAC and CGT-to-CAT), consistent with the known underrepresentation of CG (a.k.a. CpG) dinucleotides in HIV-1, likely due to methylation or ZAP [36, 37]. Because CpG-containing trinucleotide contexts are underrepresented, normalizing mutation counts by their availability increases their relative mutation rates and brings these mutation types into the higher-ranked mutation contexts despite their lower absolute mutation counts.

### NMF Resolves Two Mutational Signatures from Raw Counts and Three after Motif Normalization

The analysis of raw and motif-normalized mutation counts suggested the presence of multiple distinct mutational patterns. However, examination of individual mutation contexts alone cannot determine whether these patterns represent separable underlying mutational processes or quantify their contributions to individual viral sequences. We therefore applied non-negative matrix factorization (NMF) to deconvolve the composite mutation profiles into distinct mutational signatures (Supplementary Fig S3).

We first applied NMF directly to the raw mutation count matrix. To determine the appropriate number of mutational signatures, we evaluated candidate solutions across multiple signatures using both the average silhouette score, which measures the stability and separation of the extracted signature clusters across repeated NMF runs, and the Frobenius reconstruction error, which quantifies how accurately each model reconstructs the original mutation matrix (Supplementary Fig S4A) [26]. Considering these two criteria jointly, a two-component model representing two distinct mutational signatures was the best solution. Both signatures were dominated by the A3-associated G-to-A mutations, but in distinct sequence contexts: one enriched for GA-to-AA mutations, consistent with the sequence-context preferences of A3D, A3F, and A3H, and a second enriched for GG-to-AG mutations, consistent with A3G activity (Fig. 2A) [20, 18, 38, 39, 40, 16, 41]. Examination of the per-sequence NMF weights revealed a near-orthogonal relationship between the two mutational signatures (Fig. 2B), with sequences showing a high contribution from one signature generally exhibiting little contribution from the other. This separation between the GG-to-AG (by A3G) and GA-to-AA (by A3D/F/H) mutational patterns is consistent with previous observations, which suggested that a mechanism prevents A3G and other A3 enzymes from co-mutating the same HIV-1 genome [18].

**Fig. 2.**
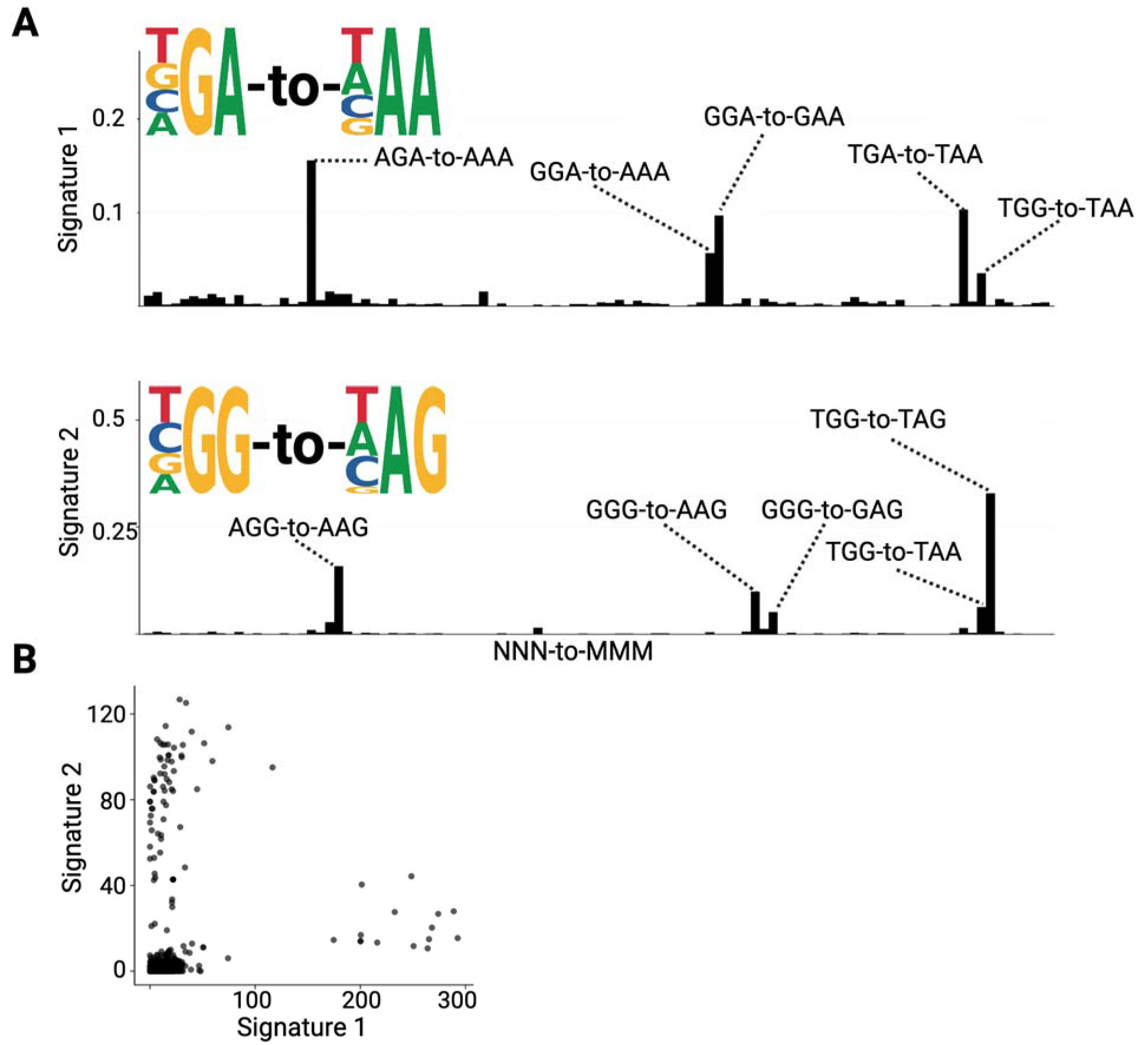
HIV-1 mutational signature deconvolution from raw mutation counts. (A) Bar plots display the relative contributions of two mutational signatures extracted by NMF. Weighted sequence logo plots define the characteristic trinucleotide substitution contexts of each signature: Signature 1 (GA-to-AA associated with A3D/F/H) and Signature 2 (GG-to-AG associated with A3G); (B) Scatter plots of pairwise signature weights demonstrate mutual exclusivity. Each dot represents an individual sequence.

We next applied the same NMF framework on mutation counts normalized by trinucleotide motif counts. Model selection was repeated independently for the motif-normalized count matrix using the same average silhouette and Frobenius reconstruction-error criteria (Supplementary Fig. S4B). In contrast to the analysis of raw mutation counts, the analysis of normalized counts showed a three-signature model as the optimum solution. Two of the three signatures recapitulated the A3-associated signatures identified in the raw mutation count analysis (Fig. 3A). Signatures 1 and 2 were characterized by G-to-A mutations within GA dinucleotides (A3D/F/H signature) and GG dinucleotides (A3G signature), respectively. Signature 3 was dominated by CGT-to-CAT substitutions. This pattern does not correspond to the canonical sequence-context preferences of known A3 family members and therefore represents a distinct, non-A3-associated mutational signature. Importantly, when considering the reverse-complement sequence, CGT-to-CAT is equivalent to ACG-to-ATG, representing a C-to-T substitution within a CpG dinucleotide. Thus, this signature may reflect methylation-associated mutagenesis, in which methylated cytosine (5-methylcytosine) within CpG sites undergoes spontaneous deamination to thymine [36, 42].

**Fig 3.**
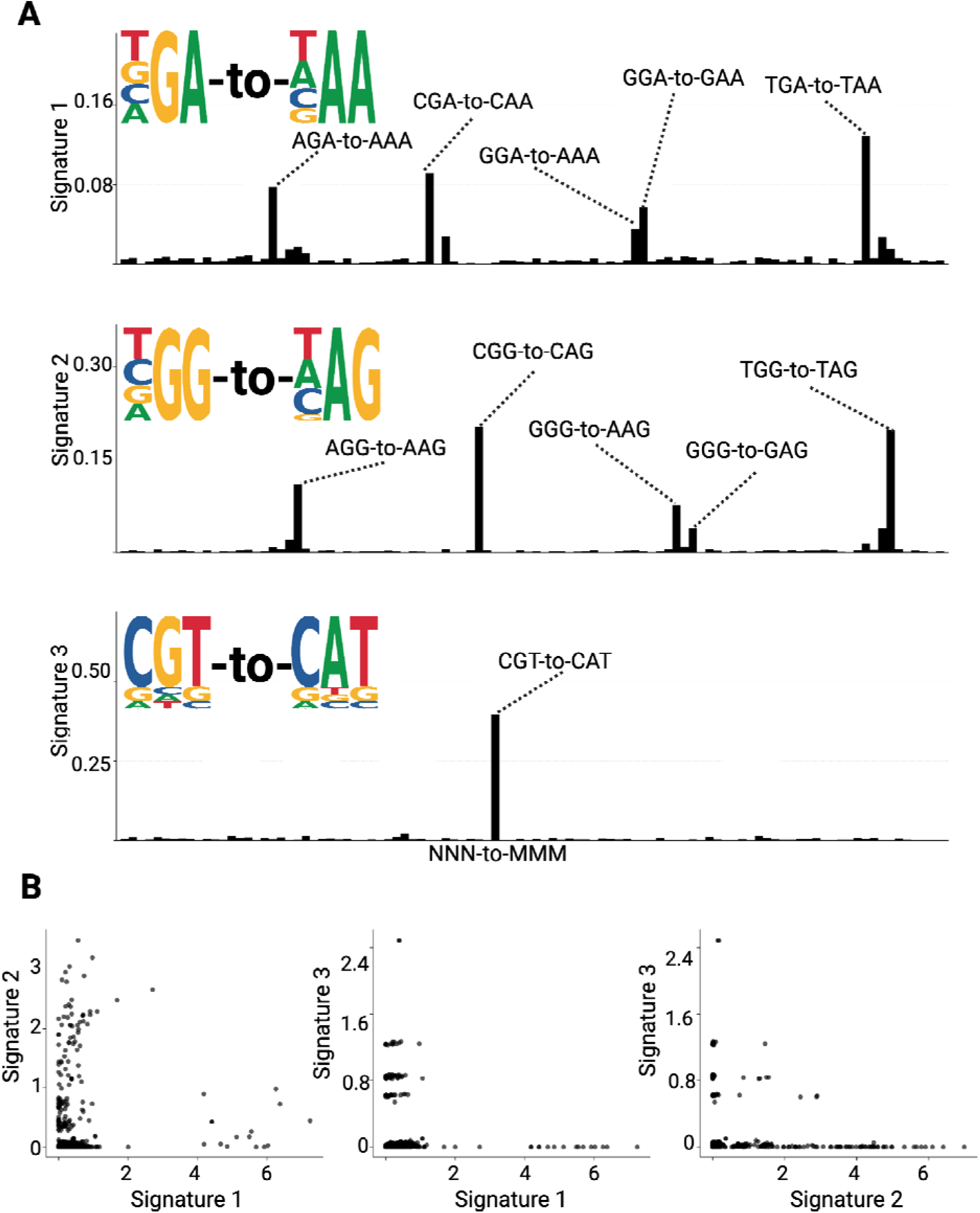
HIV-1 mutational signature deconvolution from normalized mutation counts. (A) Ba plots display the relative contributions of three principal mutational signatures extracted by NMF. Weighted sequence logo plots define the characteristic trinucleotide substitution contexts of each signature: Signature 1 (GA-to-AA associated with A3D/F/H), Signature 2 (GG-to-AG associated with A3G), and Signature 3 (CGT-to-CAT likely associated with methylation). (B) Scatter plots of pairwise signature weights demonstrate mutual exclusivity. Each dot represents an individual sequence.

The emergence of the CGT-to-CAT-dominated signature after normalization is consistent with the pattern observed in Fig. 1B. Although CGT-to-CAT mutations occur at relatively low absolute frequency and are therefore masked by the much greater abundance of A3-associated mutations in the raw data, their relative contribution increases after normalization by the number of CGT motifs in the HIV-1 sequences. Because CpG-containing motifs are underrepresented in HIV-1 genomes, this normalization increases the relative weight of CGT-to-CAT mutations and allows this lower-frequency mutational pattern to be resolved as a distinct signature.

Pairwise scatter plots of signature weights showed an approximately orthogonal pattern with limited simultaneous high contributions from multiple signatures within the same sequence (Fig. 3B). Most sequences were predominantly shaped by one signature, with relatively limited overlap between signature pairs. This pattern is consistent with limited co-occurrence of the underlying mutational processes within individual HIV-1 genomes.

Analysis of raw mutation counts showed that Signature 1 (GA-to-AA, associated with A3D/F/H) accounted for 33.8% of all mutations, while Signature 2 (GG-to-AG, associated with A3G) accounted for 16.5%. Together, these two A3-associated signatures accounted for 50.3% of all mutations (Fig. 4A). Analysis of the normalized mutation data showed that Signature 1 accounted for 29.6% of the total mutational signal, Signature 2 for 16.6%, and Signature 3 (CGT-to-CAT, likely associated with methylation) for 7.9% (Fig. 4B). The two A3-associated signatures together accounted for 46.3% of the normalized mutational signal. Thus, A3- associated mutagenesis consistently accounted for approximately half of the mutational signal in both the raw and normalized mutation count datasets.

**Fig. 4.**
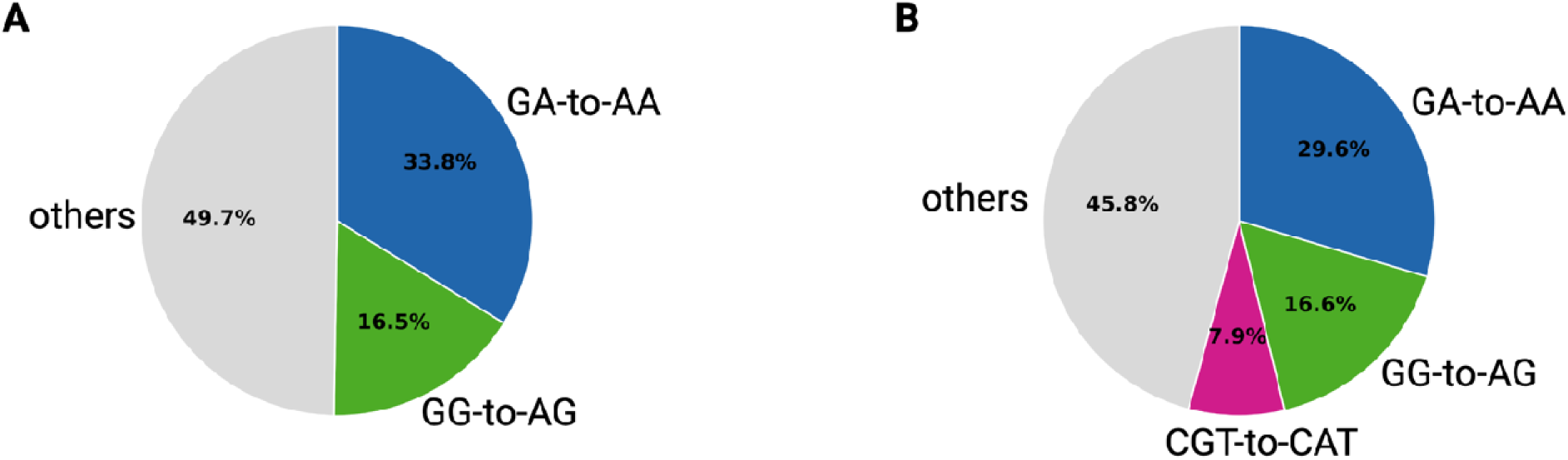
Contribution of mutational processes to the HIV-1 mutational landscape. (A) Analysis of the raw mutation counts showed that the A3-associated GA-to-AA (Signature 1) and GG-to-AG (Signature 2) accounted for 33.8% and 16.5% of the observed mutational signal, respectively, together representing 50.3% of the observed matrix. (B) Analysis of the normalized mutation counts showed that Signature 1, Signature 2, and Signature 3 (CGT-to-CAT) accounted for 29.6%, 16.6%, and 7.9%, respectively. Gray sectors indicate the remaining mutational signals not identified by NMF as having a specific pattern.

### Mutational Signatures Are Donor-Specific but Not Tissue-Specific

The NMF-derived weights of the three mutational signatures across all 1,926 sequences, ordered by donor and anatomical compartment, are shown in Fig 5.

**Fig 5.**
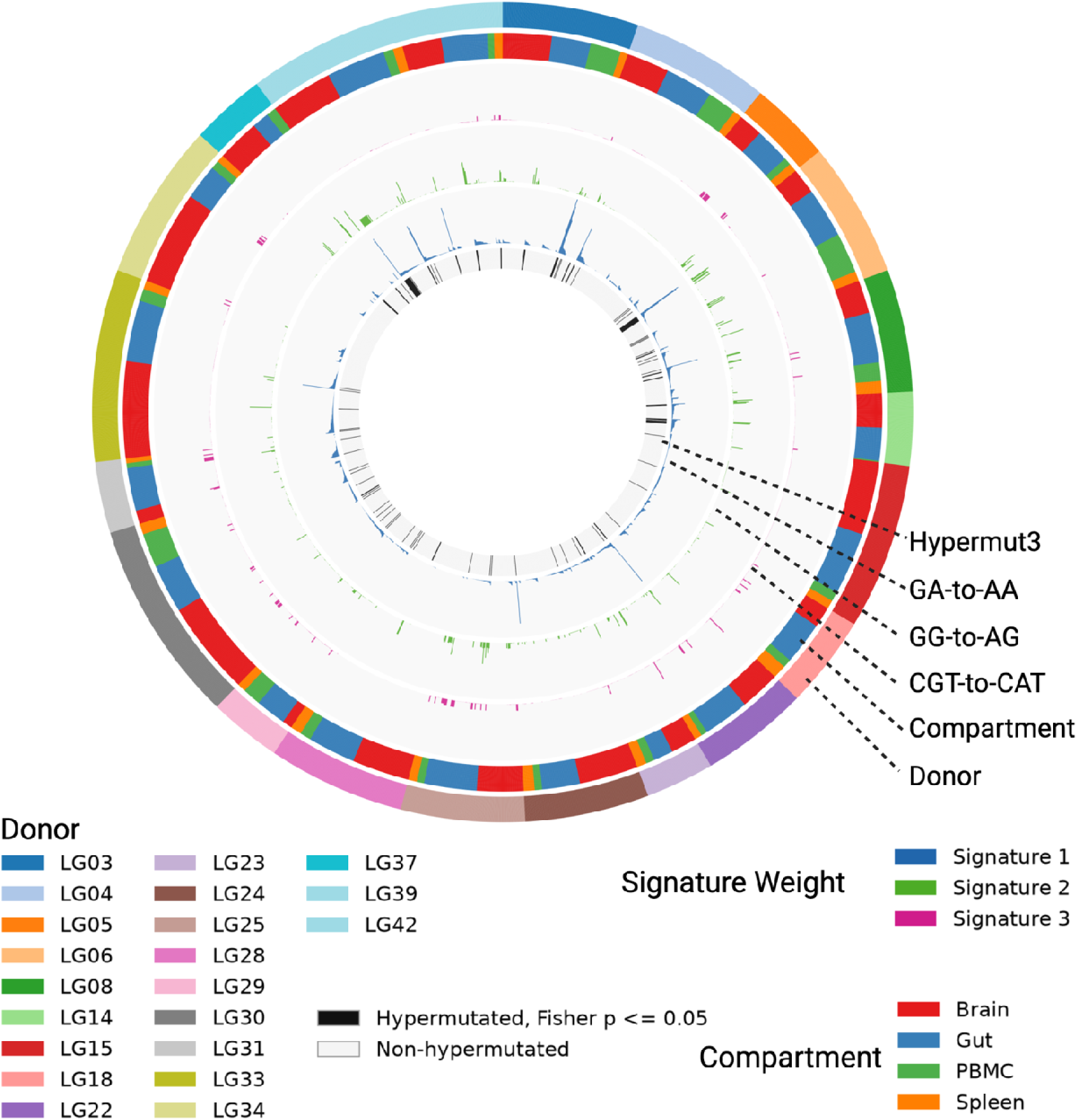
Mutational signature weights across donors and anatomical compartments. Circos plot showing signature weights for 1,926 sequences ordered by donor and anatomical compartment. From inner to outer, the tracks indicate hypermutation status, Signature 1 (GA-to-AA), Signature 2 (GG-to-AG), Signature 3 (CGT-to-CAT), anatomical compartment, and donor. The three signature tracks are displayed using a common scale.

To determine whether the three mutational signatures vary across donors and compartments, we applied a permutation-based two-way ANOVA (5,000 permutations) for each NMF-derived signature, with donor identity and compartment as main effects and their interaction as an additional term (Table 1). The analysis revealed a highly significant donor effect across all three mutational signatures (F =4.44, 5.74 and 7.94 for GA-to-AA, GG-to-AG, and CGT-to-CAT, respectively; *p = 0.0002* for all three). In contrast, the main effect of compartment was significant only for the CGT-to-CAT signature (F = 4.92, *p = 0.0104*), although the *p* value indicated only modest statistical significance, whereas no significant compartment effect was detected for GA-to-AA (F = 2.27, *p = 0.1312*) or GG-to-AG (F = 0.42, *p = 0.8602*). A significant donor × compartment interaction was likewise observed only for CGT-to-CAT (F = 2.26, *p = 0.0026*), suggesting that the effect of anatomical compartment on this signature’s weight was donor-dependent, whereas no such interaction was evident for GA-to-AA (F = 1.37, *p = 0.1498*) or GG-to-AG (F = 1.31, *p = 0.1378*).

**Table 1.** Permutation ANOVA results for mutational signature weights. Two-factor permutation ANOVA (Freedman–Lane method, 5000 permutation) was performed separately for each of the three NMF-derived mutational signatures, with donor identity, compartment, and their interactions included. Empirical p-values represent the proportion of permuted F-statistics equaling or exceeding the observed F value. Donor identity significantly influenced all three signatures. Compartment and donor-by-compartment interaction was significant only for CGT- to-CAT, indicating that compartmentalization of this signature is donor-dependent.

|  | GA-to-AA | GG-to-AG | CGT-to-CAT |
| --- | --- | --- | --- |
| Donor | $F = 4.44$<br>$p = 0.0002$ | $F = 5.74$<br>$p = 0.0002$ | $F = 7.94$<br>$p = 0.0002$ |
| Compartment | $F = 2.27$<br>$p = 0.1312$ | $F = 0.42$<br>$p = 0.8602$ | $F = 4.92$<br>$p = 0.0104$ |
| Donor × Compartment | $F = 1.37$<br>$p = 0.1498$ | $F = 1.31$<br>$p = 0.1378$ | $F = 2.26$<br>$p = 0.0026$ |

#### Post-hoc Analysis – GA-to-AA Signature

For the GA-to-AA signature, donor-level permutation without compartment stratification was applied. Significant pairwise differences were attributable mainly to one donor, LG04, which differed significantly from LG28 and LG34 (*p_adj = 0.002–0.01*,**), LG03, LG05, LG15, LG25, and LG29 (*p_adj = 0.013–0.047*,*). The additional pair that demonstrated a significant difference was LG28 and LG39 (mean difference = 0.0371, *p_adj =0.018*, *).

#### Post-hoc Analysis – GG-to-AG Signature

Since only the donor main effect was significant for the GG-to-AG signature, post-hoc analysis was performed across all compartments without stratification. Significant pairwise differences were driven primarily by two donors. Donor LG25 (mean = 0.0367) showed significantly elevated weights compared to LG05, LG34 (*p_adj = 0.008*, **); also versus LG18 and LG30 at lower significance levels (*p_adj = 0.025–0.037*, *). Donor LG06 (mean = 0.0472) also showed elevated weights relative to LG05 and LG34 (*p_adj = 0.008*, ***), LG03, LG04, LG15, LG18, LG24, LG28, LG30, LG33, (*p_adj = 0.002–0.007*, **), LG22, LG29 and LG31 (*p_adj = 0.011–0.029*, *). A significant difference in GG-to-AG signature weights was also observed between donors LG34 and LG39 (mean difference = 0.0314, *p_adj* =0.049, *).

#### Post-hoc Analysis – CGT-to-CAT Signature

Given the significant interaction term for CGT-to-CAT, Post-hoc pairwise comparisons were conducted both within each compartment (across donors) and within each donor (across compartments). Within each compartment, significant differences were confined to PBMC and gut tissues; no significant donor pairwise differences were observed in spleen or brain after MaxT correction [48]. In the PBMC, the effect was dominated by a single donor: Donor LG05 exhibited a markedly elevated CGT-to-CAT weight (mean = 0.0903) relative to virtually all other donors, with significant pairwise differences observed against LG03, LG04, LG23, LG25, LG28, LG34, (*p_adj = 0.007–0.0095*, **), LG06, LG08, LG15, LG18, LG22, LG24, LG29, LG30, LG33, LG39 (*p_adj = 0.011–0.037*, *). No other pairwise comparison in PBMC reached significance.

In the gut, only two donors exhibited pronounced effects. Donor LG25 showed the highest mean CGT-to-CAT weight (mean = 0.0565) and differed significantly from many donors, including LG03, LG04, LG06, LG15, LG23, LG24, LG28, LG30, LG33, LG39 (*p_adj = 0.0028–0.005*, **) and LG42 (*p_adj = 0.0126*, *). Additionally, Donor LG31 (mean = 0.0444) differed significantly from several donors with lower weights including LG03, LG04, LG06, LG23, LG24, LG28, LG30, LG33, LG39 (*p_adj = 0.029–0.049*, *).

Conversely, within each donor stratum, pairwise compartment comparisons showed significant differences between tissue types in four of the donors. In donor LG42, significant differences were observed between brain and spleen (mean difference = 0.0405, *p_adj = 0.021*, *), as well as between gut and spleen (mean difference =0.0345, *p_adj =0.048*, *). In donor LG33, brain differed significantly from spleen (mean difference =0.0022, *p_adj =0.014*, *). Donor LG15 showed significant differences between PBMC and brain (mean difference = 0.0012, *p_adj =0.001*,***) and between brain and spleen (mean difference = 0.0008, *p_adj =0.038*, *). Finally, in donor LG05, PBMC and spleen differed significantly (mean difference = 0.0895, *p_adj =0.020*, *).

### Cross-Compartment Sharing of Sequences Hypermutated by A3 enzymes

To investigate whether HIV-1 sequences hypermutated by A3 enzymes were confined to individual anatomical compartments or shared across tissues, we first confirmed their hypermutated status using the standard Hypermut3 program [43]. With a p ≤ 0.05 threshold we identified 123 hypermutated sequences (Gut: 55/741 = 7%; Brain: 50/805 = 6%; PBMC: 11/237 = 5%; Spleen: 7/143 = 5%; Fig 3A). Within each donor, we then assessed how similar hypermutated sequences were between different anatomical compartments using their overlapping sequence regions.

To rigorously define cross-compartment sharing of hypermutated sequences, we evaluated pairwise sequence similarity both before and after masking A3-associated mutations. Specifically, A3-associated G-to-A changes were reverted to G in each hypermutated sequence, and pairwise similarity was recalculated. This masking ensures that sequences identified as highly similar are not considered similar simply because they share A3-associated mutations but also share the remaining sequence variation. Masking increased or left unchanged the similarity of every evaluated pair (mean similarity before masking = 0.88; after masking = 0.93), supporting the interpretation that the high similarity among hypermutated sequences across tissues reflects shared ancestry.

To further distinguish genuine cross-compartment sharing from general within-donor relatedness, we required the masked similarity between the two members of each candidate hypermutated pair to be greater than or equal to the similarity of each member to every other non-hypermutated sequence from the same donor. Thus, both members of the candidate pair had to be more similar to each other than to any other non-hypermutated sequence in that donor. Pairs that did not meet this criterion were excluded because their apparent similarity could reflect broader within-donor phylogenetic relatedness rather than a specific shared clonal origin. Using this conservative approach, we identified 12 cross-compartment pairs across three donors (LG06, LG04, and LG39), including 11 brain–gut pairs and one gut–PBMC pair (Fig. 3B). No confirmed pairs involved the spleen. These data indicate that the same hypermutated provirus can be present in multiple anatomical compartments as a result of dissemination of infected cell clones.

### A3G-associated GG-to-AG is the dominant hypermutation signature

We examined per-sequence NMF signature weights for all 123 hypermutated sequences identified using the predefined Fisher’s exact-test threshold of *P* ≤ 0.05 (gut, *n* = 55; brain, *n* = 50; PBMC, *n* = 11; and spleen, *n* = 7). Overall, the A3G-associated GG-to-AG signature (Signature 2) had the highest median weight (median = 1.1491), followed by the A3D/F/H- associated GA-to-AA signature (Signature 1; median = 0.3750), whereas the CGT-to-CAT signature (Signature 3) had a median weight of 0.0000 (Fig. 7A). However, substantial sequence-level heterogeneity in signature composition was evident within the anatomical compartments (Fig. 7B).

**Fig 6.**
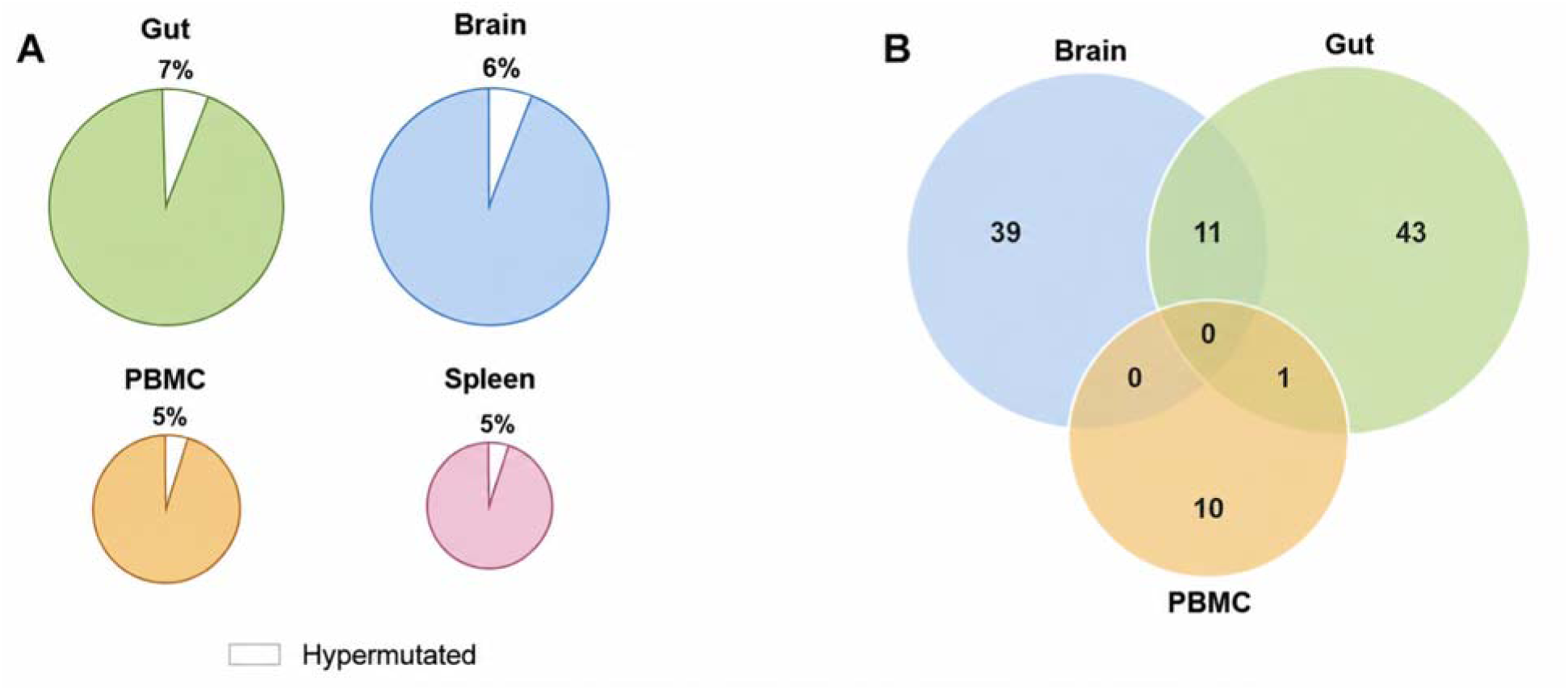
Cross-compartment sharing of hypermutated sequences. A) Pie charts showing the percentages of hypermutated sequences in each tissue, aggregated across donors: Gut, 55/741 (7.4%); Brain, 50/805 (6.2%); PBMC, 11/237 (4.6%); and Spleen, 7/143 (4.9%). The percentages were broadly comparable across the four compartments. (B) Non-area-proportional Venn diagram showing hypermutated sequences shared between anatomical compartments. Eleven sequences were shared between brain and gut, one was shared between gut and PBMC, and none were shared between brain and PBMC or across all compartments. No hypermutated sequence was shared between spleen and other compartments.

**Fig 7.**
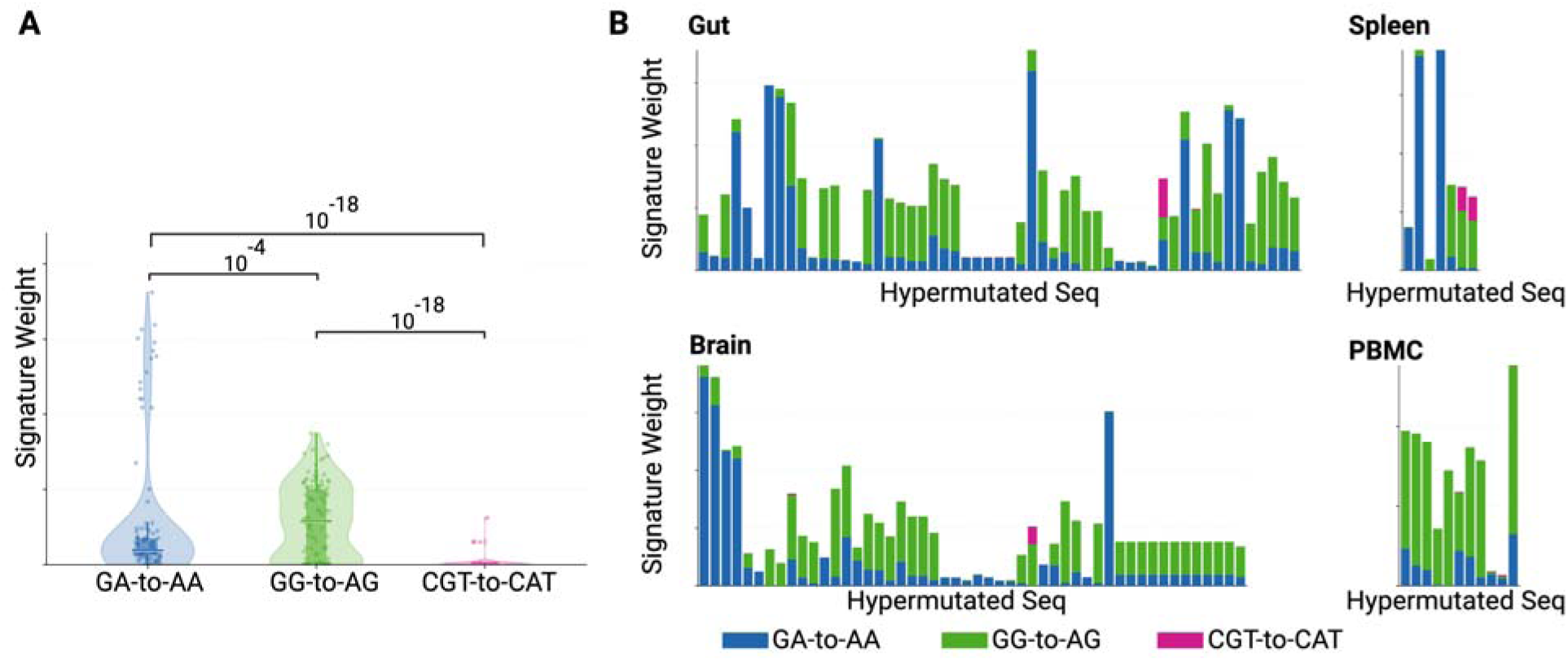
Proportion of mutational signatures in hypermutated HIV-1 sequences. **A.** Violin plots showing the distribution of signature weights across all hypermutated sequences. Embedded boxes show the interquartile range (IQR); horizontal line indicates the median; dots represent individual sequences. **B.** Bar plots showing the relative contributions of mutational signatures in individual hypermutated sequences grouped by anatomical compartment. Each bar represents a single hypermutated sequence.

Shapiro–Wilk tests indicated departures from normality for all three signature-weight distributions (GA-to-AA: *W* = 0.563, *p* = 1.7 × 10⁻¹□; GG-to-AG: *W* = 0.910, *p* = 5.3 × 10⁻□ ; and CGT-to-CAT: *W* = 0.187, *p* = 4.3 × 10⁻²³). Because all three signature weights were measured in the same sequences, differences among signatures were evaluated using a Friedman repeated-measures test. The signature weights differed significantly overall, χ²_F(2) = 127.412, *p* = 2.152 × 10⁻²□ . Paired Wilcoxon signed-rank tests showed that GG-to-AG weights were higher than GA-to-AA weights (*P_adj_* = 8.318 × 10⁻□) and CGT-to-CAT weights (*P_adj_* = 1.076 × 10⁻¹□). GA-to-AA weights were also higher than CGT-to-CAT weights (*P_adj_*= 6.971 × 10⁻¹□).

Collectively, these findings demonstrate that in this dataset A3G is the primary source of HIV-1 hypermutation. This was particularly evident in brain (p=?) and PBMC (p=?). It is important to note that while A3G-mediated GG-to-AG is the dominant signature in hypermutated HIV-1 sequences, when all HIV-1 sequences are analyzed, A3D/F/H-mediated GA-to-AA signature becomes dominant. It is known that all APOBEC3 enzymes can contribute to both extensive hypermutation and lower-level mutagenesis of HIV-1, but their relative contributions differ. A3G is more strongly associated with extensive hypermutation of individual viral genomes, whereas A3D/F/H show a greater contribution to lower-level mutagenesis across the broader viral population. Thus, A3G activity is more prominently reflected among highly hypermutated genomes, while A3D/F/H-associated mutations may affect a larger proportion of viral genomes and thereby contribute more broadly to HIV-1 sequence diversification and evolution [44].

## DISCUSSION

In this study, we applied a mutational signature deconvolution technique to extract the signatures of mutational processes in 1,926 HIV-1 *env* DNA sequences obtained from brain, gut, spleen, and peripheral blood across 21 Last Gift donors. This analysis revealed three mutational signatures, two corresponding to A3-induced GG-to-AG and GA-to-AA changes (46% of all mutations), and a third previously undescribed signature characterized by CGT-to-CAT substitutions (8% of all mutations). The remaining 46% of mutations did not exhibit a specific pattern.

The identification of two A3-associated signatures indicates that A3 enzymes are major contributors to viral genetic variation within the HIV-1 reservoir. Because A3 enzymes are expressed in a cell-type-specific manner [47, 48] and anatomical compartments comprise diverse mixtures immune and stromal cell types, we anticipated that tissues enriched for different cellular reservoirs would exhibit correspondingly distinct HIV-1 mutational signatures. Contrary to our initial hypothesis, the mutational landscape was not shaped by anatomical compartment-level factors and was instead dominated by donor-level effects.

Two scenarios may explain this lack of compartment specificity. First, CD4⁺ T cells may constitute the dominant reservoir of A3-hypermutated HIV-1 genomes across all sampled compartments, including the CNS, thereby resulting in a broadly similar mutational signature landscape across compartments. These cells could include both tissue-resident CD4⁺ T cells and clonally expanded populations that traffic between anatomical sites. Consistent with the latter interpretation, we detected identical hypermutated HIV-1 sequences shared across anatomical compartments, supporting clonal expansion and dissemination of HIV-1 infected cells across anatomical sites. However, not all hypermutated sequences were shared across compartments, suggesting that a subset of infected resident cells remains locally restricted, potentially reflecting tissue-resident CD4⁺ T cell or macrophage populations or compartment-specific persistence. Together, these findings align with growing evidence that clonal expansion of infected cells is a major mechanism of HIV-1 reservoir maintenance and that individual clones can disseminate across tissues [49, 55, 45, 48, 36].

Second, ART-mediated suppression of active viral replication may freeze the mutational landscape at a state reflecting earlier, pre-treatment dynamics rather than the current tissue-specific environment. From a therapeutic perspective, these findings underscore that reservoir elimination strategies must account for both the disseminated and tissue-resident components of viral persistence.

The CGT-to-CAT signature is of particular interest. On the complementary DNA strand, this mutation corresponds to ACG-to-ATG, a canonical CpG methylation-induced deamination that accumulates in the human genome over time. Indeed, methylation-induced deamination represents one of the most ubiquitous mutational processes in human cancer and is catalogued in the COSMIC database as the “aging” signature (SBS1). This signature is characterized by four major CpG-derived C-to-T peaks, with ACG-to-ATG being the most abundant [26, 27].

Critically, in a previous study investigating the relationship between CpG methylation rate and sequence context, we found that the ACG (complementary CGT) trinucleotide had the highest methylation rate among CpG-containing motifs and was also the most underrepresented in the HIV genome [44]. Together, these findings suggest that the HIV-1 mutational signature CGT-to-CAT may reflect CpG methylation–associated deamination, although direct experimental evidence for HIV proviral methylation in these samples is lacking.

Notably, the CGT-to-CAT (ACG-to-ATG) signature was the only signature for which the donor-by-compartment interaction was significant, suggesting that HIV-1 CpG methylation is modulated by compartment-specific factors that themselves vary across individuals.

Several limitations of this study should be acknowledged. First, sampling depth varied across donors and tissues, with the spleen being particularly underrepresented. This imbalance may have limited statistical power to detect compartment-specific effects and precluded meaningful analysis of the spleen as an independent anatomical site. Second, our analysis was restricted to the *env* gene, and the burden of mutational signatures may differ in other genomic regions. Third, although single-genome sequencing minimizes recombination and resampling artifacts, low-frequency PCR or sequencing errors cannot be fully excluded. Fourth, the cross-sectional, post-mortem design precludes analysis of temporal dynamics, providing only a snapshot of the cumulative mutational landscape at the time of death.

In conclusion, our systematic approach enabled high-resolution dissection of HIV-1 mutational signatures across anatomical compartments. The extraction of two A3 patterns and a novel CpG methylation signature illustrates the power of NMF-based deconvolution applied to viral sequence data. The strong donor-driven variation across all mutational signatures, the unexpected absence of robust tissue specificity, and observations consistent with cross-tissue dissemination of infected cell clones collectively reshape our understanding of how host factors and reservoir dynamics influence HIV-1 evolution *in vivo*.

Given that all three signatures, particularly the major A3-associated signatures, are strongly donor-specific, and that A3-induced mutations are known to contribute to drug resistance and immune evasion, future studies should prioritize identifying the host genetic factors underlying these inter-individual differences in viral mutagenesis and determining how they shape differential responses to therapy. Integrating host transcriptomic and epigenomic data with viral mutational signatures, extending the analytical pipeline to additional viral genes and larger cohorts, and experimentally investigating CpG-associated mutational processes will be essential to fully elucidate the complex interplay between host biology and viral evolution across anatomical reservoirs.

## MATERIALS AND METHODS

### Study Cohort and Sample Collection

The Last Gift cohort comprises people living with HIV who consented to participation in a rapid autopsy research protocol, enabling the collection of tissues from multiple anatomical sites shortly after death. For the present study, tissues were obtained from 21 donors and included PBMC, spleen, 5 gut regions, and up to 10 CNS regions per donor. The study was conducted with approval from the relevant institutional review boards, and all participants provided informed consent prior to enrollment. The Last Gift and CNTN studies were approved by the UCSD Human Research Protections Program (IRB#s 160563, 171024). All participants provided written informed consent for antemortem data collection and post-mortem rapid research autopsy. All procedures adhere to the Declaration of Helsinki.

### Single-Genome Amplification and Sequencing

HIV-1 envelope (*env*) DNA sequences were obtained by single-genome amplification (SGA) as described previously [1].

### Mutation Calling Pipeline

To account for the possibility that sequences within a given donor originated from more than one genetically distinct HIV-1 lineage, we implemented a lineage-aware mutation calling approach based on within-donor sequence clustering.

Sequences within each donor were first aligned to a preliminary donor-level consensus sequence. Because A3-driven hypermutation introduces dense clusters of G-to-A substitutions that could otherwise dominate the clustering step and cause hypermutated sequences to segregate artifactually into singleton or spurious clusters, all G-to-A substitutions relative to the donor consensus were provisionally reverted to G prior to clustering.

Clustering features were derived from these masked (reverted) sequences as follows. The frequency of each observed substitution at each alignment position was calculated across all sequences from a given donor, and substitutions occurring in more than 15% of that donor’s sequences at a given position were retained as informative features. Each retained (position, substitution) pair was encoded as a binary feature (1 = present, 0 = absent), yielding a binary feature vector for every sequence.

Pairwise Hamming distances between these binary feature vectors were used to cluster sequences within each donor using DBSCAN [50], evaluated across a range of epsilon (neighborhood radius) values. To select an appropriate epsilon for each donor, the total number of mutations that would be called across all sequences in that donor, computed relative to the resulting cluster-specific consensus sequences, was calculated for a series of epsilon values scanned from large to small. This yielded, for each donor, a curve of total mutation count as a function of epsilon; the epsilon value at which the first pronounced drop in total mutation count was observed was selected as the threshold for that donor, reflecting the point at which clustering resolution began to more accurately separate distinct viral lineages.

Following initial DBSCAN clustering, a post-processing step was applied to address small or noise-classified clusters. Clusters comprising fewer than 5 sequences were iteratively merged with their most similar neighboring cluster, defined as the cluster pair with the smallest average pairwise distance across all member sequences, until no cluster contained fewer than 5 sequences. Sequences classified as noise by DBSCAN were then assigned to their most similar cluster using the same average pairwise distance criterion.

For each resulting within-donor cluster, presumed to represent a distinct or closely related viral lineage, a cluster-specific consensus sequence was generated using the original (unmasked) sequences. Mutations for each sequence were called relative to this cluster-specific consensus rather than a single donor-level consensus, thereby accounting for lineage-level viral diversity within each donor. Mutations were identified within their trinucleotide context (the mutated base plus its immediate 5′ and 3′ flanking bases), yielding mutation calls of the form MMM-to-NNN (e.g., AGG-to-AAG).

To ensure unbiased comparisons across sequences of varying length and composition, mutation counts for each sequence were normalized by the frequency of the corresponding trinucleotide motif within the region overlapping the reference. Because motif counts must be obtained solely from the portion of the consensus that overlaps with each sequence, trinucleotide motifs were quantified only within this shared overlapping region. The normalized values were assembled into a mutation matrix of dimensions 1,926 sequences by 1077 trinucleotide mutation types (Fig S1A). As expected, the matrix was highly sparse, because many mutation types, particularly those involving multiple adjacent substitutions, occur infrequently.

### Non-negative Matrix Factorization

Non-negative Matrix Factorization (NMF) [26] was applied to motif-normalized mutation matrices to extract mutational signatures. Raw mutation counts were first subjected to Poisson resampling, independently for each replicate, and the resulting counts were normalized by the corresponding trinucleotide counts. For each candidate number of components (k=2–5), 1,000 independent Poisson-resampled NMF replicates were generated. NMF was implemented using scikit-learn with random initialization (init=’random’), replicate-specific random seeds, coordinate-descent optimization (solver=’cd’), and a maximum of 2,000 iterations.

For each replicate, NMF decomposed the normalized mutation matrix (X) into an exposure matrix (W) (sequences × components) and a signature-profile matrix (H) (components × mutation types), such that (X ≈ WH). The signature profiles obtained across all replicates for a given (k) were pooled and clustered using K-means with (*n_clusters_ = k*). Prior to clustering, each individual signature profile was independently L2-normalized so that clustering reflected profile shape rather than the arbitrary scale of the NMF solution. Cluster quality and signature stability were assessed using cosine-distance silhouette scores and the mean within-cluster cosine similarity.

To construct consensus mutational signatures, each signature profile within a cluster was independently normalized to sum to one before averaging. The resulting cluster centroid was subsequently renormalized to sum to one, yielding a relative mutational profile for each consensus signature. Candidate values of (k) were evaluated jointly using the average Frobenius reconstruction error across replicate NMF runs and the stability and separation of the resulting signature clusters.

After selection of the final signature set, signature exposures were re-estimated on the original, non-resampled motif-normalized mutation matrix while holding the final consensus signature profiles fixed. Non-negative least-squares (NNLS) regression was applied independently to each sequence to obtain the final exposure coefficients. Relative signature contributions for visualization were additionally calculated by normalizing the exposure coefficients within each sequence to sum to one. Final mutational signatures were visualized as bar plots showing the relative contributions of mutation types within each signature.

### Statistical Analysis

To quantify the relative contributions of donor identity and anatomical compartment to variation in NMF-derived signature weights, we performed a permutation-based two-way ANOVA for each signature independently (GA-to-AA, GG-to-AG, and (CGT-to-CAT). Donor and compartment were included as main effects, along with their interaction term, using 5,000 permutations [51, 52].

A parametric ANOVA was not appropriate because the signature weights exhibited highly right-skewed, zero-inflated distributions (GA-to-AA: 21.1%; GG-to-AG: 39.6% zeros; CGT-to-CAT: 25.9%;), violating normality assumptions. In addition, the dataset was substantially unbalanced: donors contributed unequal numbers of sequences across compartments, and two donors (LG14, LG37) lacked sequences in at least one anatomical compartment and were therefore excluded, yielding a final dataset of 1,813 sequences. Permutation ANOVA avoids assumptions of normality and balanced group sizes by generating empirical null distributions through random resampling of the observed data. For each main effect, the null distribution was generated using a restricted permutation scheme. To test the donor effect, donor labels were permuted within each compartment while compartment labels remained fixed, ensuring that only donor-level structure was disrupted [52]. The F-statistic was recalculated for each of the 5,000 permutations, and the permutation p - value was defined as the proportion of permuted F - statistics greater than or equal to the observed statistic. Similarly, to test the compartment effect, compartment labels were permuted within each donor while donor labels remained fixed.

Testing the interaction term required a different approach because simple label permutation would distort the main effects. We therefore applied the Freedman–Lane procedure, which preserves the marginal effects of both factors [53]. A reduced model containing only the main effects was first fitted, and the resulting residuals, representing variation unexplained by donor or compartment, were randomly permuted. These permuted residuals were added back to the fitted values of the reduced model, and the full model (including the interaction) was re-estimated to compute a permuted interaction F - statistic. This approach provides valid Type I error control under non-normality and unbalanced designs.

Post -hoc analyses were then performed based on the pattern of significant effects. When only the donor main effect was significant, pairwise donor comparisons were conducted using donor -level permutations without compartment stratification. The test statistic was the difference in group means expressed as a T -statistic. To control the family - wise error rate across all pairwise tests, p - values were adjusted using the MaxT procedure [54, 55] in which the maximum absolute test statistic across all comparisons is recorded at each permutation iteration. Adjusted p values were computed as the proportion of iterations in which this maximum statistic exceeded the observed statistic for a given comparison.

When the donor × compartment interaction was significant, pairwise donor comparisons were performed within each compartment separately, reflecting the fact that donor effects varied across anatomical sites. Conversely, pairwise compartment comparisons were performed within each donor separately, using compartment-level permutations without donor stratification, to assess whether the effect of anatomical compartment on signature weight varied across individual donors; p-values for these within-donor compartment comparisons were likewise adjusted using the MaxT procedure.

### Hypermutation Detection

Sequences were tested for evidence of A3-mediated hypermutation using Hypermut3 [43]. For each within-donor cluster, identified as described above, the cluster-specific consensus sequence was used as the reference sequence for hypermutation testing, rather than a single donor-level or tissue-level consensus, so that hypermutation was assessed relative to the most closely related viral lineage for each sequence. A p-value threshold of 0.05 was applied to classify sequences as hypermutated. The proportion of hypermutated sequences relative to total sequences was compared across tissues.

### Cross-Compartment Sequence Comparison

To determine whether hypermutated HIV-1 sequences were shared across anatomical compartments, we first calculated pairwise similarity among all sequences classified as hypermutated (Hypermut3, p ≤ 0.05) within each donor. For each sequence pair, a pairwise alignment was performed, and the alignment score was computed only within the overlapping region between the two sequences; The overlapping region was defined after trimming the alignment-introduced terminal gap runs at the 5′ and 3′ ends. The alignment score was then normalized by the length of the overlap region to yield a similarity value bounded between 0 and 1.

To rigorously determine HIV-1 hypermutation sharing across anatomical compartments, we masked the A3-associated G-to-A mutations with GG and GA dinucleotides in each hypermutated sequence by reverting them to the ancestral G. Reversion was guided by within-donor sequence clustering: each hypermutated sequence was assigned to its corresponding cluster, and the cluster consensus sequence was used as a reference to identify positions representing G-to-A substitutions, which were then reverted. Pairwise similarity was recalculated among the same hypermutated sequence pairs using these masked sequences ("similarity after masking"). Pairwise similarities before and after masking are shown in Supplementary Table S2).

To further confirm that high post-masking similarity reflected genuine clonal relatedness rather than shared background variation or general phylogenetic proximity, each masked hypermutated sequence was additionally compared against all non-hypermutated sequences from the same donor. A hypermutated sequence pair was retained as a confirmed "shared" pair only if its mutual (masked) similarity was greater than or equal to its similarity to every other non-hypermutated sequence within the same donor. If any non-hypermutated sequence showed similarity to either member of the pair equal to or exceeding the pair’s mutual similarity, the pair was excluded from the shared category, since the elevated similarity could not be unambiguously attributed to a specific clonal relationship between the two hypermutated sequences. This conservative approach defines the cross-compartment sequence sharing reported in Fig 3B. The complete set of within-donor hypermutated sequence-pair comparisons, including excluded pairs, is provided in Supplementary Table S2.

## ACKNOWLEDGEMNET

We are profoundly grateful to the Last Gift donors and their families for their extraordinary generosity in contributing to HIV research. This work was supported by the Texas Developmental Center for AIDS Research (D-CFAR), R56AI174877, R01AI179465-01A1, P01 AI169609 (Leaving, Coming and Staying HIV Obligate Microenvironments, HOME), the James B. Pendleton Charitable Trust, the San Diego Center for AIDS Research (SD CFAR), an NIH-funded program (P30 AI036214), which is supported by the following NIH Institutes and Centers: *NIAID, NCI, NHLBI, NIA, NICHD, NIDA, NIDCR, NIDDK, NIMH, NIMHD, NINR, FIC, and OAR,* NIDA R01DA055491.

## SUPPLEMENTARY INFORMATION

**Supplementary Table S1.** Donor information

**Supplementary Table S2.** Similarity between hypermutated sequences

---

**Supplementary Fig S1.**
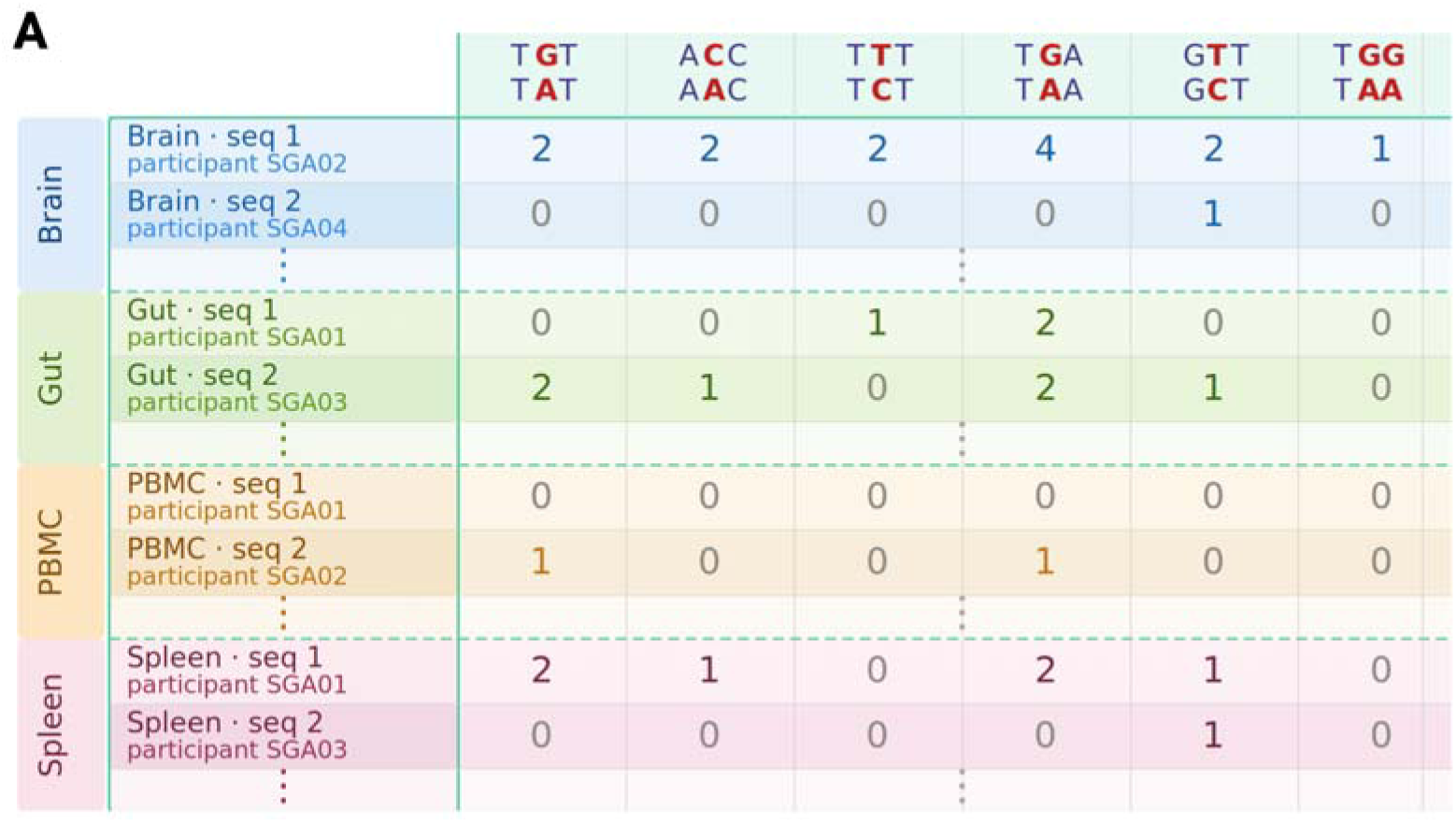
Structure of the HIV-1 trinucleotide mutation matrix. A) Each row represents a single *env* sequence and each column a trinucleotide mutation type (e.g., TGG-to-TAA). Cell values indicate mutation counts.

**Supplementary Fig S2.**
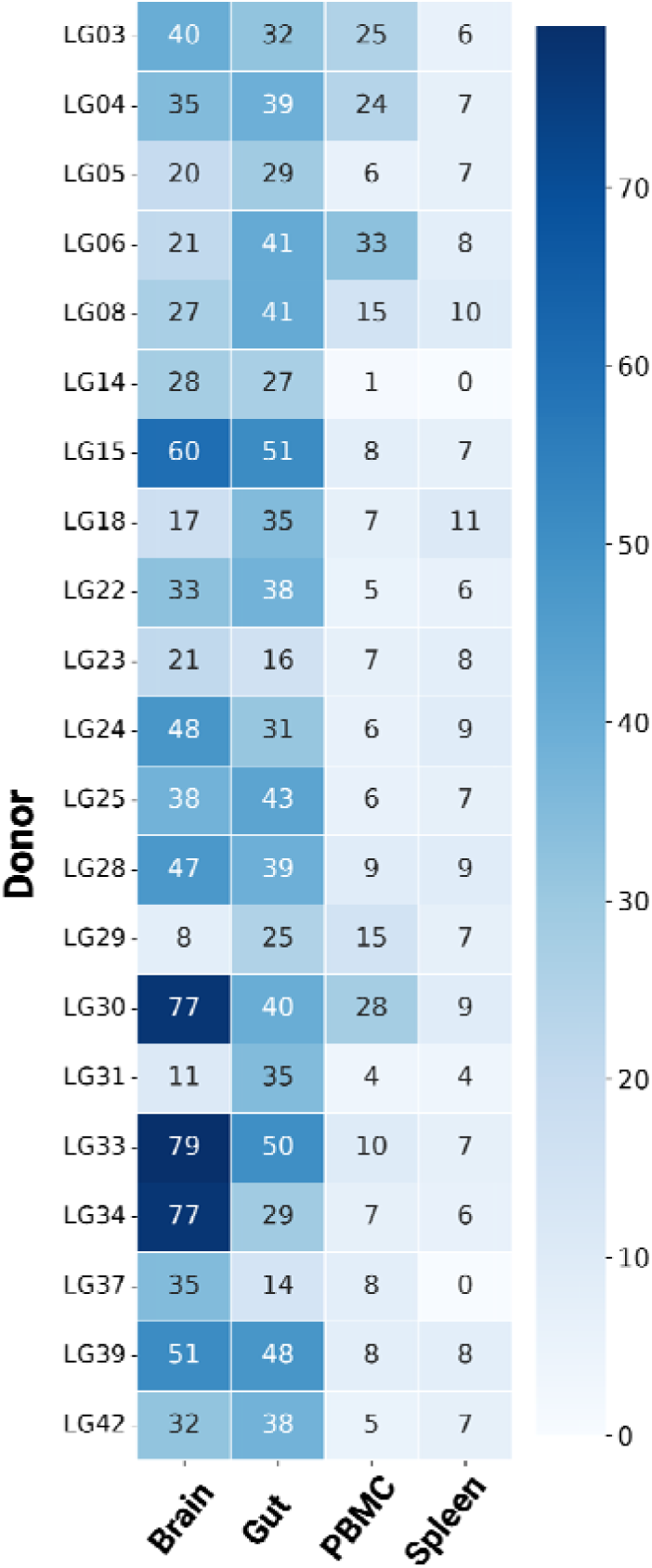
Heatmap of sequencing depth across donors and anatomical compartments. Each cell indicates the number of HIV-1 *env* sequences obtained for a given donor-tissue combination. The heatmap highlights variability in sampling depth across individuals and anatomical sites.

**Supplementary Fig S3.**
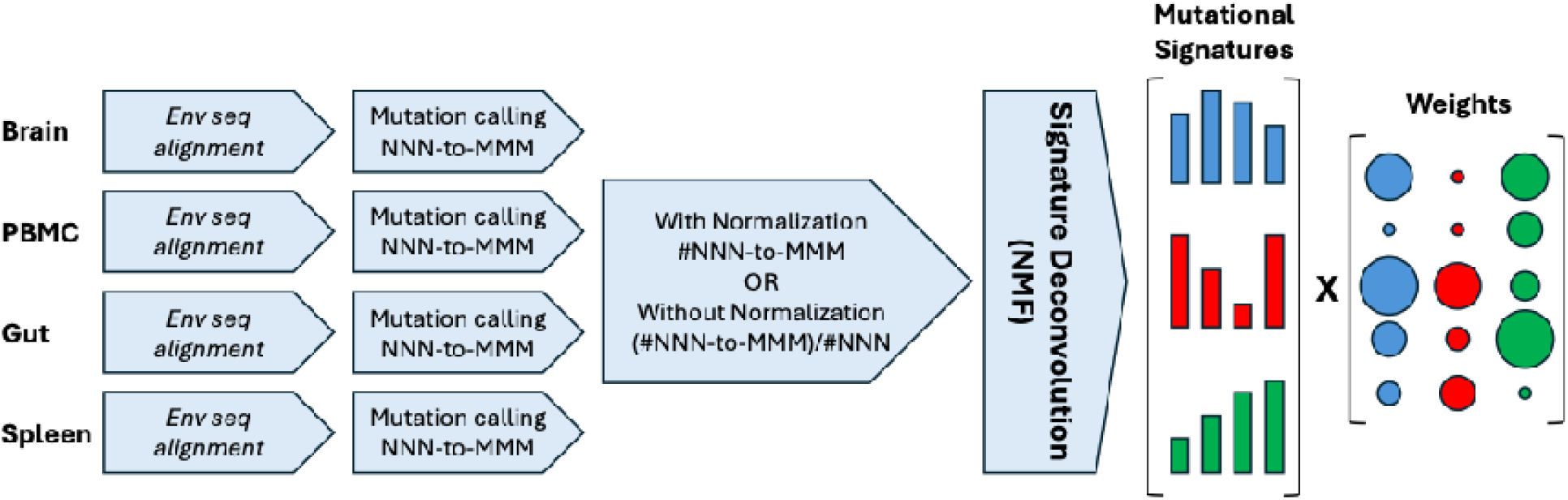
Workflow diagram of the mutation calling and NMF pipeline. The pipeline includes sequence alignment to within-site consensus, trinucleotide context extraction of mutations, normalization by genomic background frequencies of trinucleotides, and data aggregation into a mutational matrix for NMF deconvolution.

**Supplementary Fig S4.**
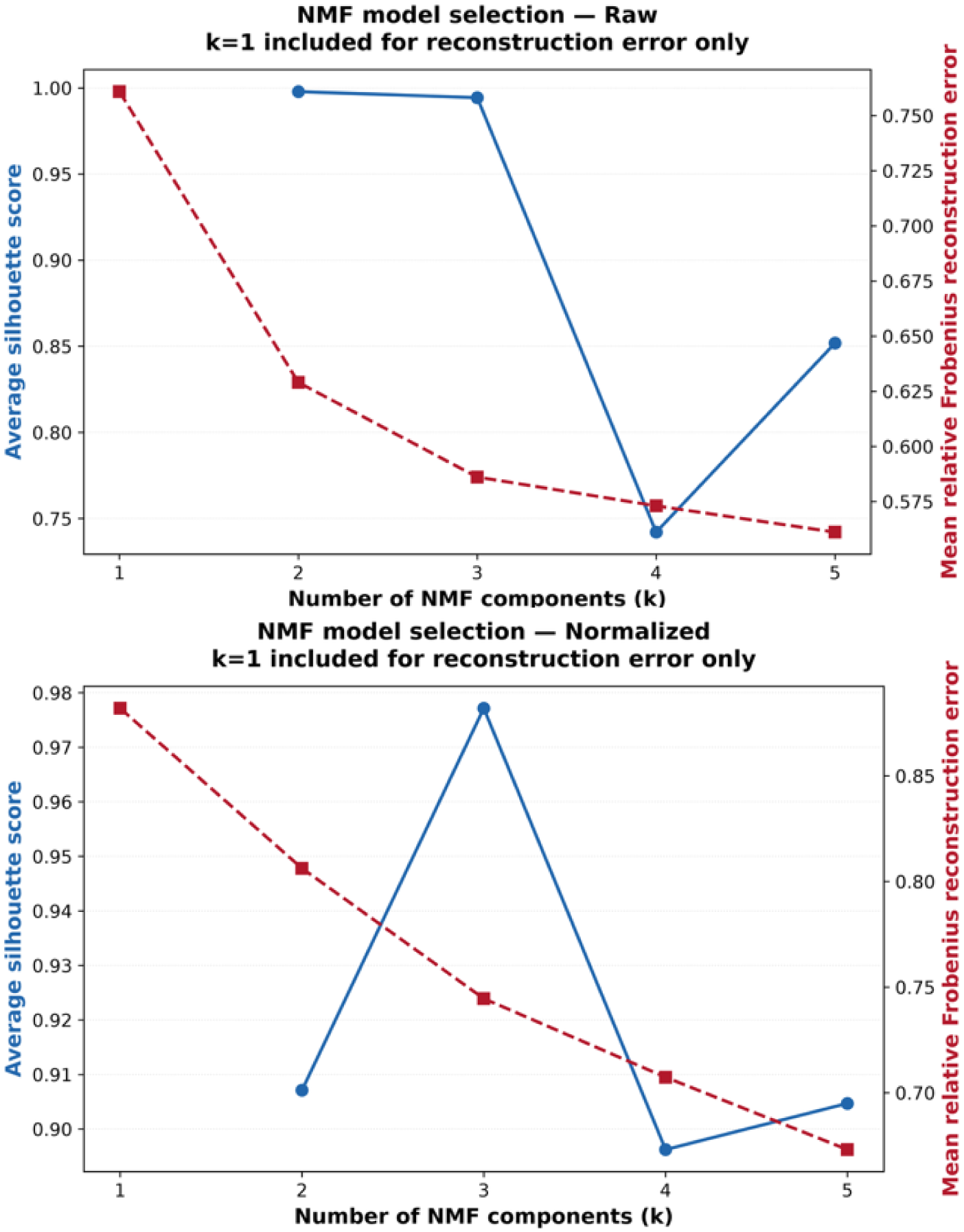
NMF model selection based on Frobenius error and Silhouette score. The average Frobenius error (blue solid line, left axis) was computed across k = 1-5 NMF components, where lower values indicate better matrix reconstruction. The Silhouette score (red dashed line, right axis) was evaluated for k = 2-5, where higher values reflect greater cluster cohesion and separation. Based on the elbow in the Frobenius error curve and the stabilization of the Silhouette score k = 2 for the raw mutation counts and k=3 for the normalized mutation counts provide an optimal balance between reconstruction accuracy and cluster quality.

